# Engineering Treg-mediated immune tolerance via *foxp3a* overexpression to evade allograft transplantation barriers in zebrafish

**DOI:** 10.1101/2025.10.15.682525

**Authors:** Junwen Zhu, Yongkang Hao, Fenghua Zhang, Xiaxia Gao, Houpeng Wang, Xiaosi Wang, Yonghua Sun

**Author notes:** Corresponding author. (X. Wang), (Y. Sun). These authors contributed equally to this work (Junwen Zhu, Yongkang Hao).

## Abstract

In mammals, regulatory T cells (Treg) cells have been utilized to enhance tolerance in organ transplantation. Although transplanting germline stem cells (GSCs) or gonadal primordia into immunodeficient zebrafish have shown to be an important technique which can bypass the juvenile phase and expedite gamete generation, it remains challenging to raise the immunodeficient fish. Here, we achieved in vivo induction of Tregs by overexpressing the key transcription factor Foxp3a, generating transgenic zebrafish with robust immune tolerance. This was characterized by a significant downregulation of T cell development and homing-related genes, accompanied by a marked upregulation of immunosuppressive factors. Using these immune-tolerant fish as hosts for subcutaneous gonadal primordium transplantation (SGPT) and intraperitoneal GSC transplantation (IGCT), we markedly accelerated germ cell maturation and efficiently established stable transgenic lines. Transcriptomic analyses demonstrated that transgenic hosts demonstrated a phenotypic profile characterized by delayed immune activation, attenuated responsiveness, and enhanced graft survival. Thus, we present a Treg-induction-based approach in fish that resolves the intrinsic conflict between high embryonic transgenesis efficiency and early lethality, while significantly improving overall transgenesis success rates. Moreover, it offers a practical alternative to immunodeficient recipients, avoiding the challenges of complex husbandry and compromised fertility.

## Introduction

Regulatory T cells (Tregs) are an immunosuppressive subset of CD25^+^CD4^+^ T lymphocytes that play an indispensable role in maintaining immune homeostasis (Sakaguchi *et al*, 2008). Their functions are implicated in diverse pathophysiological processes, including tumor immunity, autoimmune diseases, organ transplantation, and graft-versus-host disease (GvHD) (Dikiy & Rudensky, 2023; Sumida *et al*, 2024). Treg cells constitutively express high levels of co-inhibitory molecule cytotoxic T lymphocyte antigen 4 (CTLA-4), which is a key molecular target for controlling Treg-suppressive function, by competing the ligand binding with costimulation receptor CD28 on conventional T (T_conv_) cells to attenuate T cell activation (Krummel & Allison, 1995; Wing *et al*, 2008). Furthermore, Treg cells can secrete immunosuppressive cytokines such as IL-10, IL-35, and TGF-β to suppress local inflammation; simultaneously, by highly expressing the IL-2 receptor (CD25), they competitively consume IL-2 in the microenvironment, thereby inhibiting the activation and expansion of effector T cells (Dikiy & Rudensky, 2023; Sakaguchi *et al*., 2008; Vignali *et al*, 2008). Direct depletion of adaptive and natural killer immune cells through genetic editing can significantly reduce immune responses to non-self-antigens, enabling xenotransplantation of multiple tissues and organs (Ito *et al*, 2002; Yan *et al*, 2019). However, this irreversible genomic editing struggles to meet the demands for dynamic regulation within complex immune microenvironments. In contrast, Treg cell therapy demonstrates a superior risk-benefit ratio in organ transplantation and autoimmune diseases due to its physiological suppression mechanisms, cost-effective manufacturing, and dynamically adjustable properties.

In mammals, Treg cells have been utilized to enhance tolerance in organ transplantation (Wang *et al*, 2015; Yamada *et al*, 2023). Forkhead box protein P3 (Foxp3) is master regulator for the Tregs lineage, and plays crucial roles in the differentiation, maintenance, and function of Treg cells (Fontenot *et al*, 2003; Hori *et al*, 2003; Khattri *et al*, 2003; Tsuda *et al*, 2017). Specific and continuous Foxp3 expression is crucial for maintenance of the developmental immunosuppressive programs in mature peripheral Tregs (Williams & Rudensky, 2007). Two major subsets of Treg cells are identified based on their developmental origin: thymic Treg (tTreg) cells, and induced Treg (iTreg) cells (Kanamori *et al*, 2016). Currently, establishing durable immune tolerance through *in vitro* expansion and *in vivo* induction of Tregs (iTregs) has become a significant technical approach (Fontenot *et al*., 2003; Honaker *et al*, 2020; Hori *et al*., 2003; Khattri *et al*., 2003). However, research on post-transplant immune rejection and immune tolerance in fish remains limited. Recently, Foxp3 orthologs has been identified in numerous fish species (Wei *et al*, 2013; Wen *et al*, 2011; Yang *et al*, 2012; Zhang *et al*, 2011), with functional studies demonstrating that Foxp3a (but not Foxp3b) plays a critical role in Treg cell functionality (Kasheta *et al*, 2017; Li *et al*, 2020; Sugimoto *et al*, 2017). Consequently, Foxp3a appears evolutionarily conserved and represents a marker gene for teleost regulatory T cells (Hui *et al*, 2022; Kikuchi, 2020). This suggests that Foxp3a may serve as a master regulator of iTregs in teleost, positioning it as a candidate therapeutic target for achieving immunotolerance in xenotransplantation studies.

Transgenic technology and CRISPR/Cas9 gene editing (knockout/knock-in) are fundamental tools and robust strategies for investigating gene function in modern biomedicine and life sciences (Sun *et al*, 2019). However, significant trade-offs among embryonic mutation/transgenesis efficiency, embryo survival rate, and germline transmission efficiency result in a low yield of viable gene-edited or transgenic gametes. While both conventional primordial germ cell transplantation (PGCT) and the innovative induced PGCT (iPGCT) approach effectively address germline transmission challenges in gene-edited gametes, successful offspring acquisition requires completion of the full gonadal developmental cycle in zebrafish (spanning 3 mpf) (Ciruna *et al*, 2002; Wang *et al*, 2023b; Zhang *et al*, 2020). Secondly, all embryonic manipulations are performed at the blastula stage, which only allows for genetic-level detection and precludes phenotypic observation beyond this stage. However, post-blastula phenotypic assessment is crucial for determining the success of transgenesis and gene knock-in. Notably, transplantation of gonadal primordia or germline stem cells (GSCs) derived from larvae or adult fish into adult recipients can bypass the juvenile stage and accelerate gamete production (Kawasaki *et al*, 2016). Yet, immune rejection post-transplantation hinders durable engraftment, requiring host immunosuppression through genome editing or drug treatment (Montano-Loza *et al*, 2023; Yoshinaga *et al*, 2021). Immunocompromised zebrafish exhibit inherent limitations including heightened pathogen susceptibility, elevated mortality rates, growth retardation, reproductive challenges, and substantial maintenance costs for sterile husbandry (Yan *et al*., 2019). Chronic immunosuppressant administration, though alternative, induces systemic immunosuppression—significantly increasing susceptibility to opportunistic infections, chronic organ damage, and malignancy risks (Rice *et al*, 2017; Stoletov *et al*, 2007). It remains unclear whether we can achieve an *in vivo* induction of Tregs in zebrafish to increase the efficiency of GSC transplantation and gonadal primordia transplantation.

In this study, we generated and analyzed *foxp3a*-overexpressing transgenic zebrafish lines that exhibit enhanced Tregs functionality and increased immune tolerance through suppression of effector T cell activity. When applied to high-efficiency transgenic fish as subcutaneous transplantation recipients, these zebrafish accelerated the maturation of gonadal primordia in developmentally malformed embryos and promoted the maturation of transplanted GSCs in the cloacal region. Our findings demonstrate that *foxp3a* overexpression strengthens immune tolerance in zebrafish and highlights the potential of this zebrafish in vivo Treg enhancement model for surrogate reproduction.

## Results

### Generation of *foxp3a*-ubiquitous transgenic zebrafish

Intraperitoneal germline stem cell transplantation (IGCT) (Fig. S1A) and subcutaneous gonadal primordium transplantation (SGPT) (Fig. S1B) are regarded as the most promising and effective approaches to bypass the juvenile phase and expedite gamete generation (Farlora *et al*, 2014; Kawasaki *et al*., 2016). However, immune rejection precludes the survival of gonadal primordia transplanted via IGCT and SGPT in wild-type adult fish. To circumvent this dilemma, we generated a transgenic zebrafish line, *Tg(CMV:foxp3a; CMV:mCherry)* (ihb946Tg, hereafter abbreviated as *Tg(CMV:foxp3a)*), featuring systemic overexpression of foxp3a – a key transcriptional regulator of Treg cells (Fig. 1A). This strategy is designed to induce functional Treg cells in vivo, thereby fostering immune tolerance toward exogenous transplants (Hori *et al*., 2003; Khattri *et al*., 2003). To enable post-transplantation visualization of the grafts, the *Tg(CMV:foxp3a)* line was generated on the genetic background of the *Casper* transparent zebrafish strain (CZ73). As shown in Figure 2, we successfully established a transgenic zebrafish line *Tg(CMV:foxp3a)* exhibiting ubiquitous expression of *foxp3a*, with mCherry serving as a marker for rapid identification of positive fish. To validate the expression level and pattern of *foxp3a*, we performed whole-mount in situ hybridization (WISH) on F1 embryos at 4 days post-fertilization (dpf). The results demonstrated a significant increase in *foxp3a* expression levels throughout various anatomical regions of *Tg(CMV:foxp3a)* embryos, particularly in the head, visceral organs, and musculature (Fig. 1C). Furthermore, quantitative real-time RT-PCR (RT-qPCR) analysis of tissues from 2 months post-fertilization (mpf) fish— including thymus, spleen, head kidney, mid kidney, gills, liver, intestine, heart, eye, brain, skin, and muscle – revealed significantly elevated *foxp3a* expression across all examined tissues in *Tg(CMV:foxp3a)*. Strikingly, expression in the gills exhibited an upregulation of several hundred-fold (Fig. 1D). Collectively, these results demonstrate that we have successfully established a transgenic zebrafish line exhibiting ubiquitous high-level expression of *foxp3a*.

**Fig. 1.**
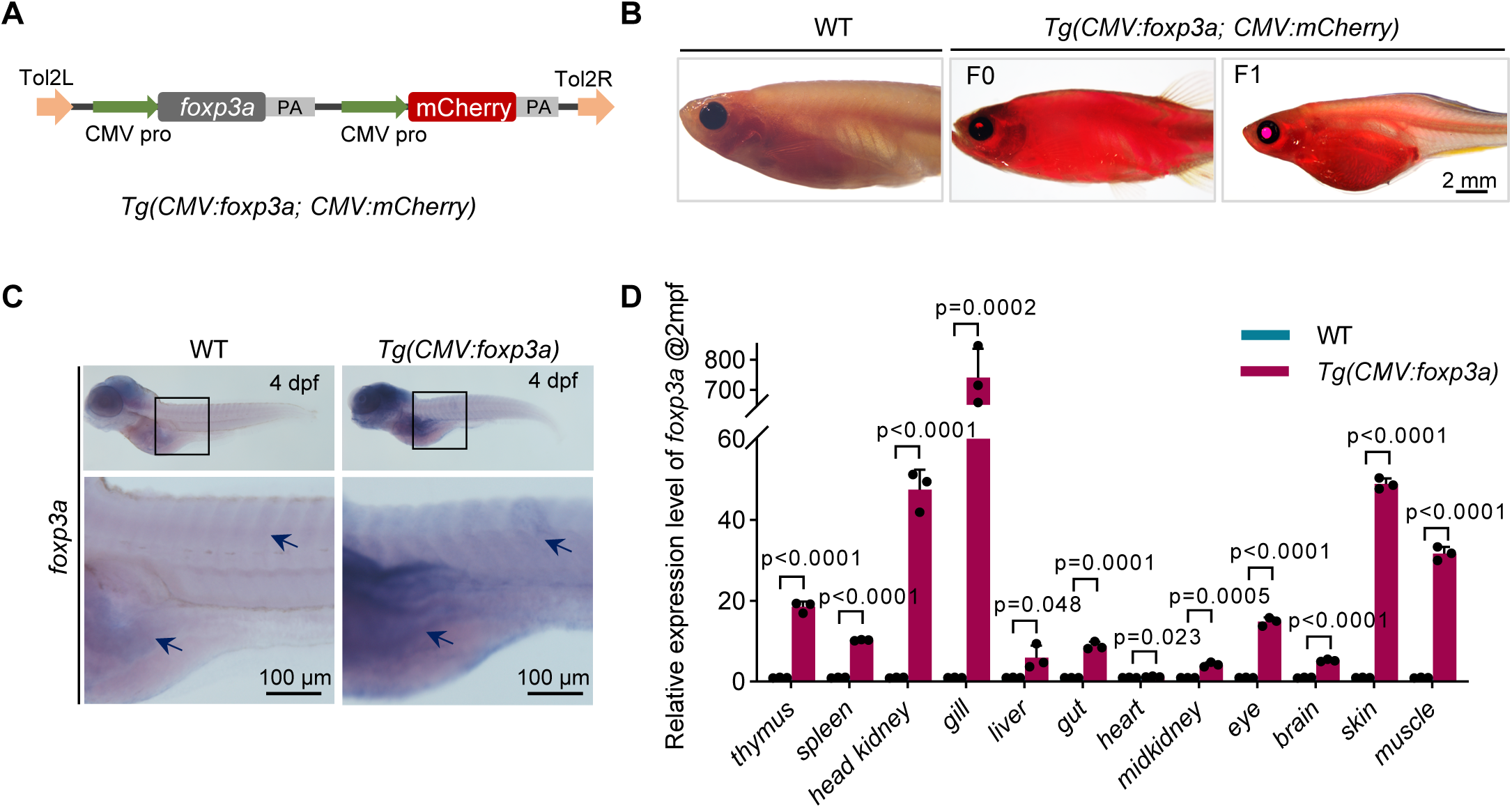
Construction of a Foxp3a-Overexpressing Transgenic Zebrafish Line. (A) Schematic of the *foxp3a* overexpression strategy. Using the Tol2 transgenic vector, the target gene *foxp3a* was inserted downstream of a *CMV* promoter followed by a SV40 poly A tail’s signal. Another *CMV* promoter drived mCherry fluorescent protein expression for fluorescence screening. F1 and R1 marked on the vector represented forward and reverse primers for PCR-based specific screening of positive fish, with one primer binding to the *foxp3a* gene and the other to the vector’s poly A tail. (B) *Casper* wild-type and fluorescent-positive *Tg(CMV:foxp3a)* transgenic fish lineages in F0 and F1 generations. (C) Brightfield images showing in situ hybridization results of *foxp3a* RNA probe in WT and *Tg(CMV:foxp3a)* transgenic fish, n=30. (D) RT-PCR analysis of *foxp3a* expression in various tissues of WT and *Tg(CMV:foxp3a)* transgenic fish at 2 mpf. t-test; *p*<0.05 indicates statistically significant difference.

**Fig. 2.**
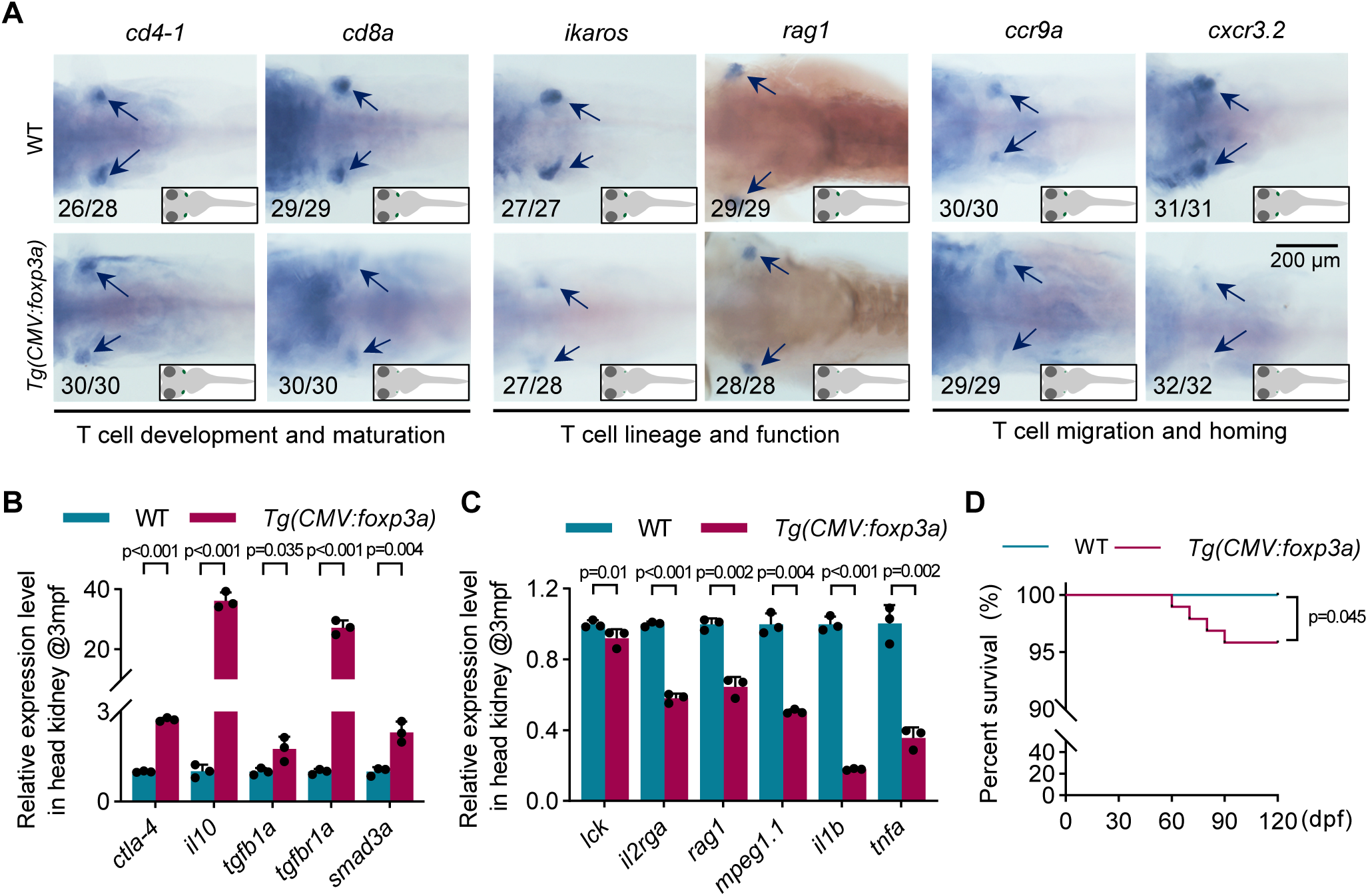
Immune characterization of *Tg(CMV:foxp3a)* transgenic fish (A) Bright-field images displaying in situ hybridization results of *cd4-1*, *cd8a*, *ikaros, rag1*, *ccr9a*, and *cxcr3.2* RNA probes in WT and *Tg(CMV:foxp3a)* transgenic fish at 4 dpf, n=30. (B) RT-PCR analysis showing significantly elevated *ctla-4*、*il10*、*tgfb1a*、*tgfb1ra* and *samd3a* expression levels in head kidney at adult, n=3, t-test, *p*<0.05 represented significant difference. (C) RT-PCR analysis showing significantly elevated *lck*, *il2rga*, *rag1*, *mpeg1.1*, *il1b*, and *tnfa* expression levels in head kidney at adult, n=3, t-test, *p*<0.05 represented significant difference. (D) Survival curves of WT, and *Tg(CMV:foxp3a)* from 4 dpf to 120 dpf, t-test, *p*<0.05 indicates statistically significant difference, n=96.

### *Tg(CMV:foxp3a)* transgenic fish demonstrate immune tolerance phenotypes

To further assess the immunocompetence of the *Tg(CMV:foxp3a)* transgenic fish, we conducted comprehensive analyses of immune cells and immune factors using WISH and RT-qPCR. CD4⁺ T cells, predominantly helper T cells (Th cells), represent a major subset of T lymphocytes. Foxp3 upregulation or overexpression drives CD4⁺ T cell differentiation toward the Treg lineage, accompanied by downregulation of CD8 and Th1/Th17-associated genes, resulting in attenuated immune responses, while immunosuppressive factors (CTLA-4, IL-10, TGF-β) are upregulated (Honaker *et al*., 2020; Sakaguchi *et al*., 2008). In the thymus of *Tg(CMV:foxp3a)*, expression of *lck* (a T cell—specific tyrosine kinase) (Kasheta *et al*., 2017) and *cd4-1* remained largely unchanged (Fig. 2A, S2A and S2B), whereas immunosuppressive factors (*cd28*, *il-10*, *tgfb1a*, *tgfbr1a*, *smad3a*) were markedly upregulated in the head kidney (Fig. 2B). Interestingly, *in vitro* Foxp3-overexpression in tilapia and zebrafish robustly enhanced transcriptional activation of anti-inflammatory cytokines TGF-β1 and IL-10, while concurrently upregulating the inhibitory receptor CTLA-4 (also known as CD28) (Kasheta *et al*., 2017; Sugimoto *et al*, 2006). These findings suggest that while T cell numbers remain stable, their functional state is altered, with an expansion of the Treg population. Conversely, in the thymus of *Tg(CMV:foxp3a)*, expression of the cytotoxic T cell factor *cd8* (Ellmeier *et al*, 2013), the master regulator of lymphocyte development *Ikaros* (also known as *ikzf1*) (Oravecz *et al*, 2015), and the key gene *rag1* required for early T and B cell development (Wienholds *et al*, 2002) was markedly downregulated (Fig. 2A), indicating impaired development and maturation of T cells, particularly CD8⁺ cells. In addition, expression of the chemokine receptor *ccr9a*, which guides T cell homing to the thymus and related tissues, and *cxcr3.2*, a key receptor for effector T cell migration, was significantly reduced (Fig. 2A), indicating impaired T cell trafficking and homing in *Tg(CMV:foxp3a)*. At the same time, RT-PCR analysis of the head kidney in adult fish revealed significant downregulation of T/B cell-related genes *il2rga* and *rag1*, macrophage-associated gene *mpeg1.1*, and proinflammatory cytokine genes *il1b* and *tnfa* in *Tg(CMV:foxp3a)*. Overall, in these *foxp3a*-overexpressing fish, the immune milieu is characterized by Treg dominance and enhanced immunosuppression, accompanied by impaired T cell development, differentiation, and trafficking. Consequently, the overall immune system displays a propensity toward immune tolerance.

Interestingly, survival curves from 4 to 120 dpf revealed that *Tg(CMV:foxp3a)* exhibited survival rates comparable to those of controls. Under standard aquaculture system conditions, they exhibited over 98% survival rate at 60 dpf and maintained above 95% survival rate at 90 dpf (Fig. 2D). Given that immune-deficient mutants often exhibit poor survival and reduced reproductive capacity under standard rearing conditions (Kawasaki *et al*., 2016; Yan *et al*., 2019), *Tg(CMV:foxp3a)*, with its immune tolerance and ease of maintenance, holds promise as a preferred recipient for transplantation.

### Allogeneic gonadal tissue can colonize and proliferate subcutaneously in *Tg(CMV:foxp3a)*

To assess the potential of immune-tolerant *Tg(CMV:foxp3a)* fish as surrogate reproductive hosts, we performed subcutaneous transplantation assays. Because high germline transmission efficiency of transgenes is often associated with increased embryonic deformity, we further carried out subcutaneous gonadal primordium transplantation (SGPT), in which tissue blocks containing only donor embryonic gonadal primordium were implanted subcutaneously into 2-mpf host fish (Fig. 3A). To facilitate unambiguous germ cell labelling, we generated *Tg(ddx4:GFP)* transgenic zebrafish. However, transgene-positive embryos frequently exhibited developmental malformations due to plasmid toxicity and failed to transmit the germline to subsequent generations. To circumvent this limitation, we surgically removed extraneous tissues— including the head, trunk, and visceral organs—from transgene-positive F0 embryos, retaining only the gonadal ridge as graft material for subcutaneous gonadal primordium transplantation (Fig. 3B). Because endogenous germ cells can compete with transplanted germ cells, recipient germ cells were ablated by injecting *dnd1* MO during embryogenesis (Zhang *et al*, 2022; Zhang *et al*., 2020). Graft colonization was monitored daily after transplantation, with photographic documentation every 7 days. Beginning at 7 days post-transplantation (dpt), grafts gradually underwent clearance by the host. Between 14 and 21 dpt, however, exogenous gonadal primordia partially engrafted in *Tg(CMV:foxp3a)* fish and initiated proliferation. By contrast, grafts in wild-type hosts exhibited rejection responses during this period, characterized by tissue disaggregation and progressive diminution of cellular fluorescence, and eventually complete disappearance (Fig. 3C). Surprisingly, by 35 dpt, *Tg(CMV:foxp3a)* hosts had successfully developed markedly enlarged grafts densely populated with vasa-positive cells (Fig. 3D). This suggests that germ cells within the grafts likely underwent rapid proliferation and differentiation. Continuous observations of the survival time of grafts in WT and *Tg(cmv:foxp3a)* indicated that the key time window for exogenous gonadal primordia immune rejection was 7-14 dpt in zebrafish (Fig. 3E). Overall, subcutaneous gonadal primordium transplantation enabled successful engraftment and sustained proliferation of allogeneic gonadal tissue in immune-tolerant *Tg(CMV:foxp3a)* fish, underscoring their potential as a valuable platform for immune tolerance and transplantation studies.

**Fig. 3.**
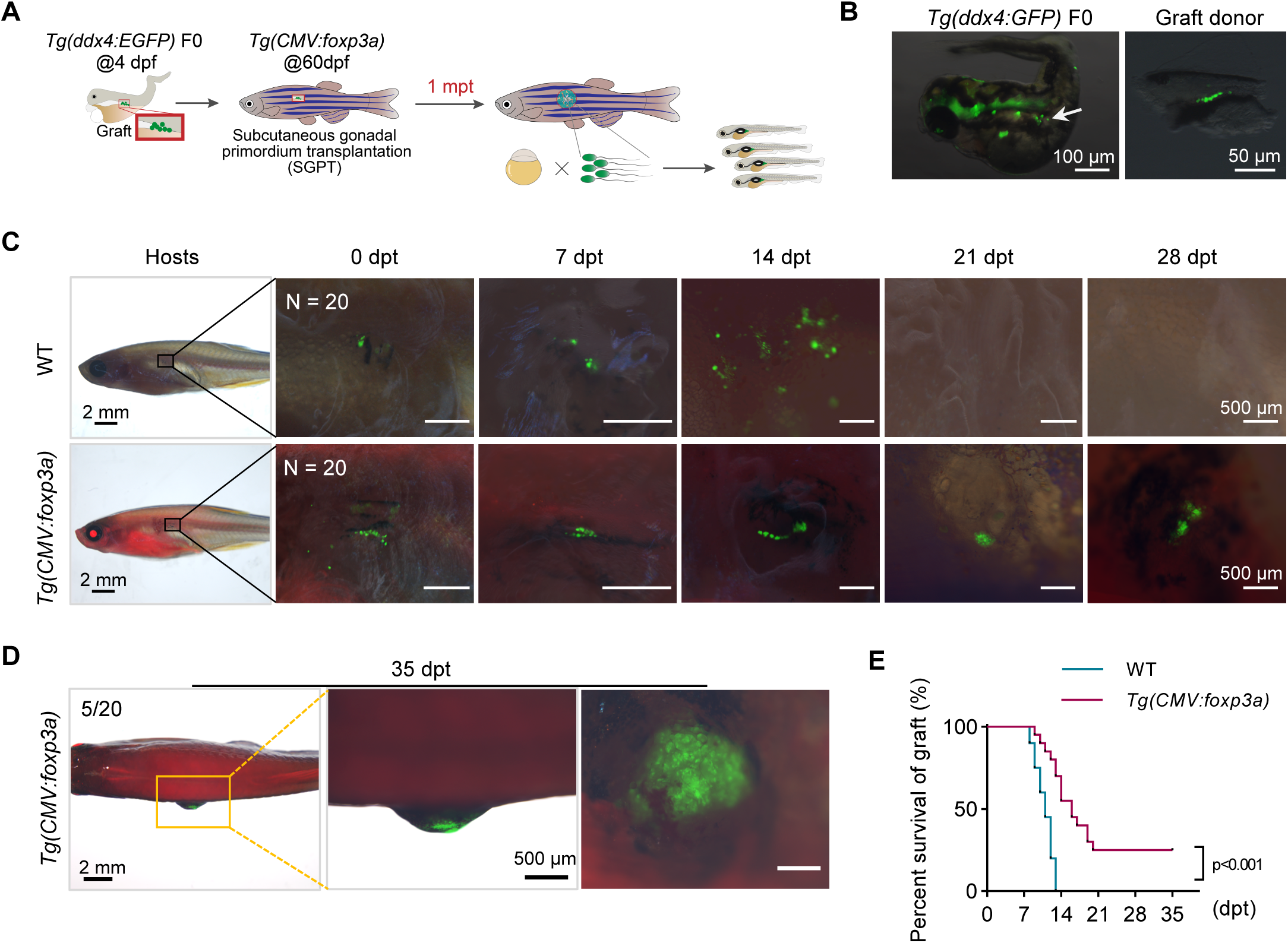
*Tg(CMV:foxp3a)* transgenic tolerant fish for gonadal gland subcutaneous transplantation (A) The flowchart of subcutaneous transplantation of gonadal primordium (4 dpf) in host fish (60 dpf) to quickly produce functional sperm. (B) At 4 dpf, the green fluorescent labeling of germ cells in *Tg(ddx4:GFP)* F0 funder donor fish and the generation of grafts derived from gonadal primordia. (C) Form 0 dpt, imaging tracking of subcutaneous grafts in WT and *Tg(CMV: foxp3a)* tolerant fish every 7 days, with green fluorescently labeled Ddx4 positive germ cells, n=20. (D) Bright-field images showing proliferating grafts, with green fluorescently labeled Ddx4 positive germ cells at 35 dpt, n=5. (E) Survival curve of grafts under the fish’s skin, t-test, *p*<0.05 indicates statistically significant difference, n=20.

### Immunological dynamics in the graft and host fish thymus post-subcutaneous transplantation

To further investigate the immune dynamics and host responses within grafts following subcutaneous gonadal primordium transplantation (SGPT) in both wild-type and *Tg(CMV:foxp3a)*, we performed RNA-seq analysis on grafts and thymic tissues at 10 dpt (initial rejection phase in wildtype) and 14 dpt (completion of wildtype rejection vs. successful grafts colonization in *Tg(CMV:foxp3a)* hosts (Fig. 4A). Given that not all *Tg(CMV:foxp3a)* hosts supported germ cell proliferation and development in SGPT (approximately 25% of hosts developed hypertrophied grafts; Figure 3D and 3E), we extended RNA-seq analysis to thymic tissues from *Tg(CMV:foxp3a)* hosts that failed to sustain graft viability. In the early stages of allogeneic transplantation, host CD8⁺ T cells typically recognize donor-derived MHC class I molecules on graft cells through their T cell receptors (TCRs), thereby eliciting acute rejection (Klawon *et al*, 2025). Strikingly, transcriptomic analyses at 10 dpt revealed significantly higher expression of *mhc1uba* and *mhc1uka* in both the thymus and grafts of *Tg(cmv:foxp3a)* hosts compared with controls (Fig. 4B and S2A). This finding suggests that reduced expression of *cd8a* and chemokine receptors in *Tg(cmv:foxp3a)* attenuates effector T cell responses, permitting sustained expression of MHC class I molecules. However, within the graft, molecules associated with innate immune activation (*tlr4ba*, *il1b*, *tnfa*, *cxcl8a*), oxidative stress (*nos2a*, *nos2b*), antigen processing (*psmb9a*), and immune checkpoint regulation (*hhla2a.1*) were markedly upregulated (Fig. S2A and S2B), consistent with a prototypical inflammatory amplification response. By contrast, in control recipients, CD8⁺ T cells rapidly respond to graft-expressed MHC class I, triggering rejection and resulting in a marked reduction of MHC class I levels (Fig. 3C, 3E and 4B). As MHC-I molecules (*mhc1uka*, *mhc1uba*) remained persistently upregulated in *Tg(cmv:foxp3a)* hosts, antigen presentation in the thymus was sustained, leading to activation of cell-mediated immune pathways associated with inflammation and rejection, such as the induction of Th17 cytokines (*il17a/f3*) (Fig. 4B). However, elevated *foxp3a* expression reinforced Treg-mediated tolerance at 10 dpt, placing the immune system in a paradoxical state of being simultaneously activated yet restrained.

**Fig. 4.**
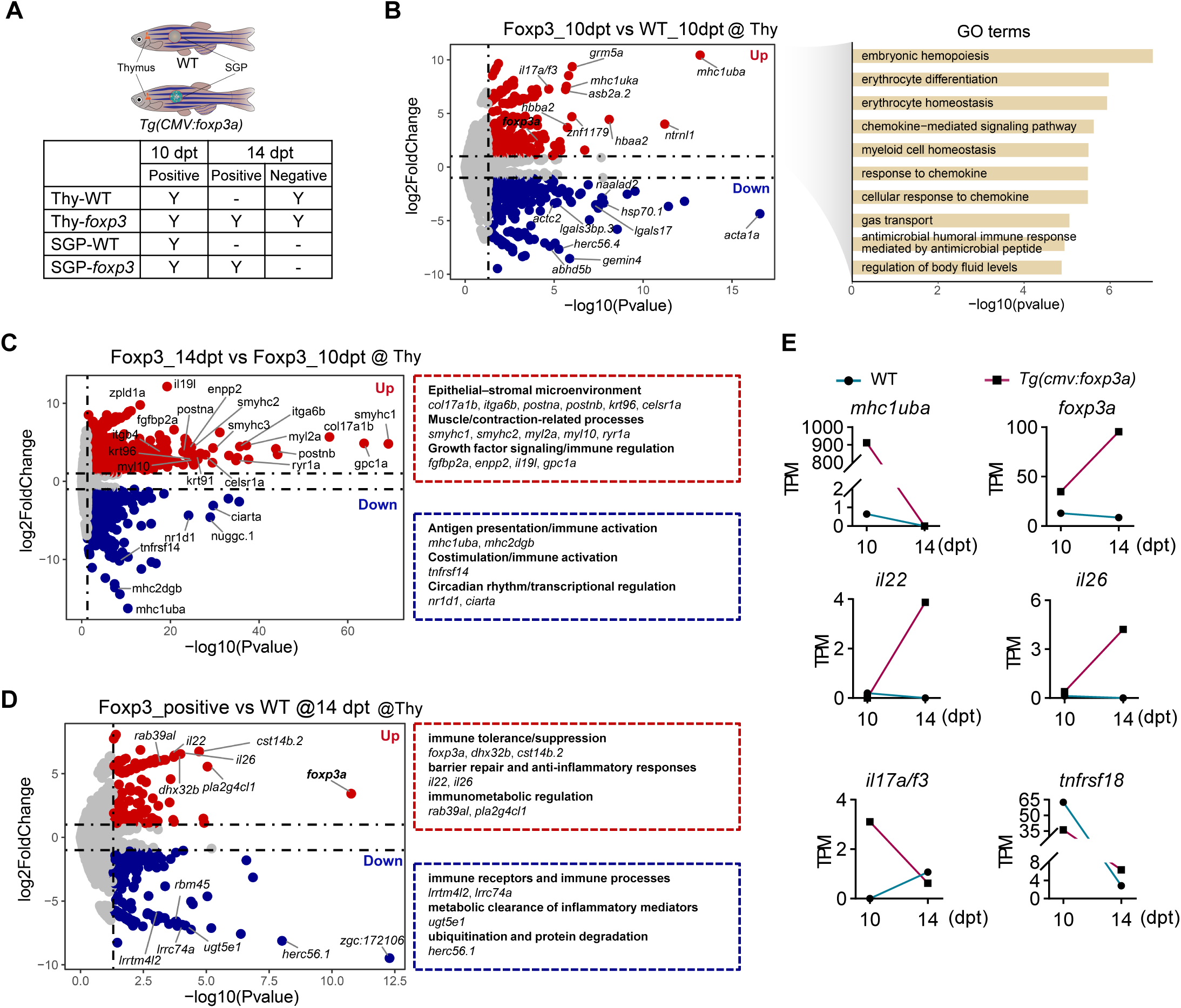
Immunological processes in the graft and host fish thymus following subcutaneous implantation (A) The flowchart of the sample for RNA-seq. (B) The Volcano Plot of WT and *Tg(CMV:foxp3a)* thymus at 10 dpt, and GO terms with up regulated signal pathway. (C) The Volcano Plot of *Tg(CMV:foxp3a)* thymus at 10 dpt and 14 dpt, and GO terms with up and down regulated signal pathway. (D) The Volcano Plot of WT and *Tg(CMV:foxp3a)* thymus at 14 dpt, and GO terms with up and down regulated signal pathway. (E) WT and *Tg(CMV:foxp3a)* thymus immune related gene (*mhc1uba*, *foxp3a*, *il22*, *il26*, *il17a/f3* and *tnfrsf18*) vary by TPM at 10 dpt and 14 dpt.

At 14 dpt, thymic transcriptomes of *Tg(cmv:foxp3a)* hosts revealed attenuated antigen presentation (*mhc1uba*, *mhc2dgb*), diminished immune activation signals (*tnfrsf14*), and reduced expression of circadian regulators (*nr1d1*, *ciarta*) (Fig. 4C), indicating a shift toward an immunosuppressive state that prevents excessive immune activation. Consistently, *foxp3a*, *il22*, and *il26* were upregulated, supporting enhanced Treg activity, barrier repair, and establishment of a tolerogenic environment. Conversely, genes linked to immune recognition and stress responses (*rbm45*, *lrrtm4l2*, *lrrc74a*, *ugt5e1*, *herc56.1*) were downregulated, while factors associated with epithelial–stromal remodeling (*postna*, *postnb*, *col17a1b*), muscle contraction (*smyhc1*, *smyhc2*, *myl2a*, *myl10*), and growth or immune modulation (*fgfbp2a*, *enpp2*, *il19l*) were upregulated (Fig. 4C), suggesting strengthened tissue remodeling and repair. Notably, graft transcriptomes at the same time point exhibited a global reduction of circadian regulators (Fig. S4C, S4D), consistent with reduced metabolic activity, dampened immune responsiveness, and improved graft survival.

Comparative analyses further revealed divergent transcriptional programs between recipients that supported versus rejected graft engraftment. Successful engraftment was characterized by upregulation of *il26*, *rsad2*, *ifi44a7*, *bmp10*, and *wnt1*, implicating pathways involved in tissue repair, antiviral defense, and developmental regulation, thereby restraining inflammation and fostering regeneration. In contrast, rejection-prone individuals showed elevated expression of epithelial stress and injury markers (*krt93*, *krt95*, *mlsl*, *vmn2r1*) together with Ca²⁺ excitability and proteolysis factors (*cacna1fb*, *prss59.1*, *dpep1*, *lgals9l5*), reflecting an epithelial-stress–driven microenvironment that promotes effector immune clearance. Taken together, *Tg(cmv:foxp3a)* hosts mounted a delayed immune response characterized by progressive accumulation of MHC class I molecules and concomitant *foxp3a* induction, thereby shifting the immune balance toward tolerance and repair, ultimately restraining excessive activation and promoting graft survival.

### Accelerated generation of donor-derived functional transgenic sperm via subcutaneous gonadal transplantation

As previously described, using *Tg(CMV:foxp3a)* hosts with SGPT enables generation of germ cell-replete grafts within just one month (Fig. 3D). To evaluate the developmental status of these germ cells, we performed immunofluorescence staining on the grafted tissues at 35 dpt, as well as on control gonadal tissues at 40 dpf (corresponding to the same developmental stage as the grafts) and 90 dpf (representing mature testes) (Fig. 5A). The fluorescent tissue portions of the transplant were excised and subjected to immunofluorescence staining using germline stem cell marker Nanos2, germ cell marker Ddx4, meiotic cell marker Sycp3, and germ cell proliferation marker Pcna (Cao *et al*, 2019; Ye *et al*, 2023). Notably, Nanos2-positive stem cells in SGPT grafts exhibited comparable abundance to controls, indicating sustained spermatogenesis capability in the transplanted gonadal primordia (Fig. S4A and S4B). Unexpectedly, the germ cell composition in the SGP was the same as that in the 90 dpf control group, containing germ cells at different stages of spermatogenesis, including mature spermatozoa (SPZ). In contrast, in the 40 dpf gonads, which are at the same developmental stage as the SGP, germ cells had only developed up to the SPC-II stage, with no spermatids formed (Fig. 5B). Furthermore, we performed detailed classification of germ cells in the grafts using two antibodies, Sycp3 and Pcna (Ye *et al*., 2023). Compared with the 40 dpf gonads, the proportion of Pcna-positive cells in the grafts was markedly lower, indicating that the proportion of mitotically active germ cells was reduced and that germ cells were likely in a more differentiated state (Fig. 5C). The Sycp3 staining results showed a reduced proportion of Sycp3-positive germ cells in the grafts, suggesting that germ cells had already completed meiosis and were mainly spermatozoa (Fig. 5D). Quantitative analysis further demonstrated that spermatozoa were significantly more abundant in the grafts than in the 40 dpf control gonads, and their germ cell composition resembled that of mature testes (Fig. 5E). Overall, these results demonstrated that SGPT accelerated the biological processes of germ cells proliferation and differentiation. Moreover, normal self-renewal of germline stem cells was maintained in proliferating gonadal tissues.

**Fig. 5.**
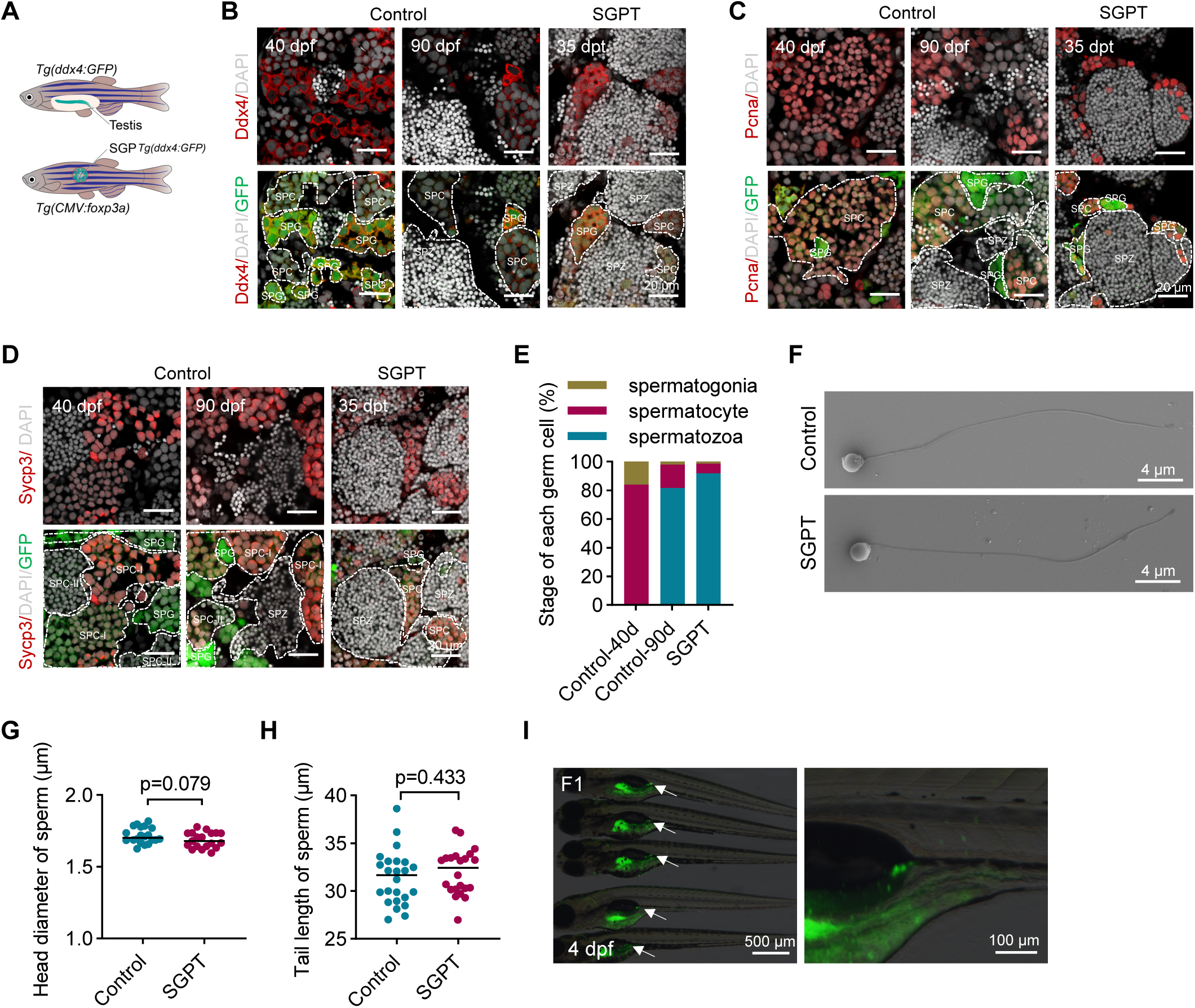
Subcutaneous transplanted gonads accelerated the biological process of proliferation and differentiation (A) The flowchart of control and SGPT testes tissue. (B-D) Immunofluorescence of tissue sections from *Tg(ddx4:GFP)* testes (40 dpf and 90 dpf) and subcutaneously grown *Tg(CMV:foxp3a)* testes (35 dpt), with Ddx4 antibody (B), Pcna antibody (C), Sycp3 antibody (D). (E) Statistical analysis of spermatogonial stem cells, spermatocytes, and sperm proportions in *Tg(ddx4:GFP)* testes (40 dpf and 90 dpf) and subcutaneously grown *Tg(CMV:foxp3a)* testes (35 dpt) sections. (F) SEM images of subcutaneous transplanted sperm and wild-type sperm and local enlargement of the head. (G) Head diameter of SRGT sperm and wild-type sperm, t-test. (H) Tail length of SRGT sperm and wild-type sperm, t-test. (I) At 4 dpf, the fluorescence of *Tg(ddx4:GFP)* zygote was displayed at the genital ridge, the place indicated by the white arrow, and the local enlargement of the genital ridge. Antibody staining appears as red fluorescence, endogenously expressed GFP as green fluorescence, and DAPI-stained nuclei as gray fluorescence. All data *P*<0.05 indicates statistically significant difference, n=3.

To verify whether these sperm were functional, we gently homogenized tissue from the enlarged grafts to release sperm. Scanning electron microscopy (SEM) analysis showed that the morphology of graft-derived sperm was comparable to that of wild-type zebrafish sperm, including equivalent head diameter and tail length (Fig. 5F-5H). Furthermore, fertilization assays with wild-type eggs demonstrated that sperm from the grafts were functional, capable of fertilizing eggs and producing normally developing embryos (Fig. 5I and S4C). Notably, although germ cell development was slightly delayed in some grafts, resulting in relatively lower fertilization rates (Fig. S4D), this germ cell “accelerated maturation” strategy can substantially reduce the time and resources required for breeding experiments. Overall, by overexpressing *foxp3a*, we established an immune-tolerant transplantation host that enables germ cells to undergo accelerated maturation.

### Germline stem cells can colonize and rapidly proliferate in the abdominal cavity of immune-tolerant fish

Intraperitoneal germ cell transplantation (IGCT) represents another promising technique for accelerating germ cell maturation. To evaluate the suitability of *Tg(CMV:foxp3a)* transgenic fish as immune-tolerant hosts for germline transplantation, we transplanted germline stem cells isolated from 3 mpf *Tg(ddx4:GFP)* donor zebrafish into the intraperitoneal cavity of 2 mpf *Tg(CMV:foxp3a)* transgenic hosts via cloacal microinjection (Fig. 6A). Donor testes were dissected and purified to obtain male germline stem cells (mGSCs). Compared to controls, Percoll density gradient centrifugation significantly enhanced the isolation and enrichment of Nanos2-positive germline stem cells (Fig. 6B and 6C). Subsequently, the isolated germ cells were transplanted via cloacal microinjection into the intraperitoneal cavity of germ cell-depleted 2 mpf host fish (Fig. 6D). At 0 dpt, GFP-labeled germ cells derived from *Tg(ddx4:GFP)* donors were observed in both wild-type and immune-tolerant hosts (Fig. 6E). At 35 dpt, all donor-derived GFP-positive cells had disappeared in the peritoneal cavity of wild-type hosts, whereas donor-derived germline stem cells were able to survive in a small proportion of immune-tolerant hosts (Fig. 6F). This indicates that germline stem cells can also rapidly proliferate and differentiate in the peritoneal cavity of 2 mpf immune-tolerant hosts. The relatively low survival rate of germline stem cells may be attributed to individual variability among *foxp3a* transgenic fish, as well as to the overall low number of transplanted germline stem cells. Intriguingly, despite initially low fertilization rates, transplantation-positive host fish spontaneously mated with wild-type females without requiring artificial fertilization assistance. More significantly, as transplanted fish matured, egg fertilization rates progressively increased to exceed 80% (Fig. 6G and 6H). This suggests progressive gonadal maturation within the intraperitoneal cavity, yielding increased production of functional sperm. Further, embryonic fluorescence imaging unambiguously confirmed these sperm originated from the donor rather than the host (Fig. 6I). Overall, germline stem cells exhibit robust colonization capacity and undergo rapid proliferative expansion within the peritoneal microenvironment of immunotolerant teleost species.

**Fig. 6.**
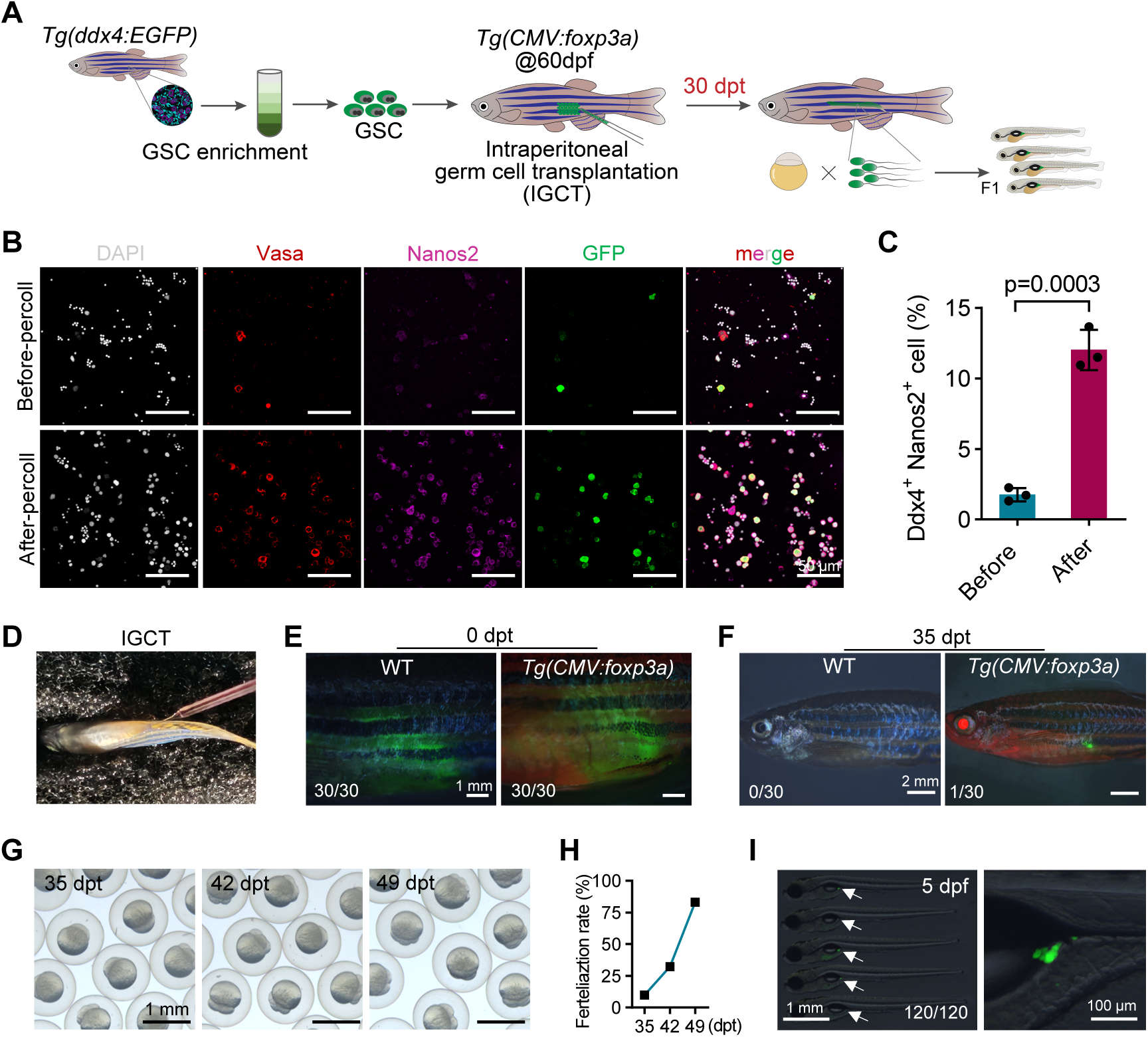
*Tg(CMV:foxp3a)* transgenic fish accelerates maturation of GSCs transplanted via the adult cloacal pore (A) Schematic diagram illustrating the rapid generation of functional sperm via cloacal transplantation of germline stem cells (GSCs) into germ cell-depleted recipient fish (60 dpf). (B-C) Immunofluorescence staining of Nanos2 and Ddx4 before and after Percoll density gradient centrifugation purification of GSCs (B), and quantification of Ddx4^+^/Nanos2^+^ double-positive cells before-and after-Percoll purification (C). t-test, *p*<0.05 indicates statistically significant difference, n=3. (D) Schematic of intragonadal germ cell transplantation (IGCT). (E-F) Representative images of germ cell colonization in WT and *Tg(CMV:foxp3a)* hosts at 0 dpt (E) and 35 dpt (F), n=30; experiments repeated four times. (G-H) Photographic documentation (G) and statistical analysis (H) of sperm production rates in positive recipient fish at 35 dpt, 42 dpt, and 49 dpt. (I) Fluorescence imaging of *Tg(ddx4:GFP)* zygotes at 4 dpf, showing GFP expression in the genital ridge (white arrows), with insets highlighting magnified views of the genital ridge.

## Discussion

In this study, we successfully generated transgenic zebrafish with immune-tolerant traits using a *foxp3a* overexpression–based induced Treg strategy. Transcriptomic and in situ hybridization analyses demonstrated that the overall fish exhibited an immune-tolerant microenvironment. This was characterized by a significant downregulation of T cell development and homing-related genes, accompanied by a marked upregulation of immunosuppressive factors. More importantly, these *foxp3a*-overexpressing zebrafish served as excellent recipients for both intraperitoneal germline stem cell transplantation (IGCT) and subcutaneous gonadal primordium transplantation (SGPT), enabling accelerated maturation of germ cells for applications such as transgenesis.

As a lineage-defining transcription factor governing Treg cell development and maintenance, Foxp3 plays a pivotal role in sustaining Treg-mediated immune homeostasis. *In vitro* and *in vivo* induction of *foxp3* expression has been well-established to upregulate Treg-mediated immune tolerance in mammalian models (Fontenot *et al*., 2003; Honaker *et al*., 2020; Hori *et al*., 2003; Khattri *et al*., 2003). Studies in fish species have yet to explore the induction of immunosuppressive T cells (iTregs) through *foxp3a* overexpression—although *foxp3a* has been demonstrated to play essential roles in regulating immune homeostasis in gonadal, intestinal and other tissues (Kasheta *et al*., 2017; Li *et al*., 2020; Sugimoto *et al*., 2017). Our study successfully generated an immune-tolerant recipient fish by overexpressing *foxp3a*. In this transgenic line, the expression of T cell development-related genes such as *cd8*, *ikaros*, and *rag1* was markedly reduced in the thymus, whereas immunosuppressive factors including *il10*, *ctla-4*, and members of the TGF-β family were significantly upregulated in the head kidney. These results suggest a notable expansion of Foxp3a-positive regulatory T cells in the transgenic fish. More importantly, using organ transplantation—the gold standard for assessing immune tolerance—we demonstrated that the *foxp3a* transgenic fish indeed possesses an immune-tolerant microenvironment capable of supporting the development of allogeneic gonadal primordia or germline stem cells. It is worth noting that in human organ transplantation, immune modification is usually performed on the donor organ (Wang *et al*, 2023a), whereas in fish, immune modification primarily targets the transplant recipient. Therefore, investigating the dynamic changes in the immune microenvironment following transplantation in fish may provide valuable complementary insights for human organ transplantation research.

In vertebrates, immunodeficient mutants are widely employed in allogeneic and xenogeneic transplantation studies. Zebrafish models with single or multiple gene knockouts have enabled successful transplantation of human tumor cells (Yan *et al*., 2019). However, such immunocompromised models exhibit elevated mortality rates and require long-term antibiotic-treated rearing systems with meticulous maintenance due to their compromised immune systems lacking partial or multiple immune cell populations (Yan *et al*., 2019). In comparison, this Treg-induced immunotolerant strain can be maintained under standard zebrafish husbandry protocols while exhibiting comparable viability to wild-type counterparts, thereby establishing a robust paradigm for conventionalized cultivation of transplantation hosts. This successful establishment of a tolerant zebrafish line addresses technical challenges associated with immune-deficient zebrafish hosts and circumvents adverse effects of immunosuppressive regimens.

The technology of PGCT, iPGCT, and GSCT can rapidly and efficiently improve the efficiency of the genome editing and germline transmission of zebrafish (Wang *et al*., 2023b; Xie *et al*, 2022; Zhang *et al*., 2022). However, there is a considerable gap between PCGT/iPGCT transplantation and the ultimate acquisition of gametes. This is because the expression of fluorescent proteins cannot be directly observed prior to transplantation, especially in gene knock-in or transgenic experiments where the target gene is expressed after the blastocyst stage. Moreover, the number of isolated and purified germline stem cells is crucial for GSCT and is closely related to the success rate of transplantation (Zhang *et al*., 2022). Our methodology incorporates a high-efficiency transgenesis platform with three innovative functionalities: (i) real-time fluorescent screening for successfully edited events, (ii) germline transmission efficiency approaching 100%, and (iii) a significant acceleration of gametogenesis timelines. This establishes a more robust and practical solution for precision genome engineering applications. Notably, our current study achieved functional sperm production exclusively through SGPT. This limitation arises because germ cell-depleted zebrafish recipients exclusively developed into males (Slanchev *et al*, 2005). Establishing an infertile female recipient may enable the rapid acquisition of donor-derived functional eggs. On the other hand, within hypertrophied grafts, we observed not only expanded germ cell populations but also developing other tissues. However, likely due to competitive constraints in the recipient microenvironment, the development of these extraneous tissues was compromised (Kawasaki *et al*., 2016). This finding suggests that employing organ-deficient recipients could enable our established immune-tolerant subcutaneous transplantation platform to facilitate rapid generation of functional organs.

In this study, we demonstrate that engineering Treg-mediated immune tolerance via *foxp3a* overexpression enables successful partial SGPT and IGCT. However, this ubiquitous-overexpression of *foxp3a* represents a broad yet unrefined approach. Future work should focus on either T cell-specific enhancement of *foxp3a* expression or epigenetic modification-mediated upregulation of *foxp3a* (Honaker *et al*., 2020; Khattri *et al*., 2003). Nevertheless, our findings represent only an initial step in host modification. Subcutaneous transplantation experiments revealed that merely 25% of tolerant recipients sustained full allograft colonization, proliferation, and differentiation. The remaining 75% demonstrated delayed but ultimately complete rejection by 21 dpf (Fig. 3E). This may be related to individual variability among *foxp3a* transgenic fish. On the other hand, differences in innate immune responses during transplantation are also an important factor influencing whether grafts can successfully engraft (Yu *et al*, 2025). Therefore, future modifications of transplantation recipients should fully take into account the role of innate immune responses. On the other hand, emerging evidence indicates that, in addition to Foxp3^+^ Treg cells, mammals also possess other immunoregulatory populations, including IL-10–producing Breg cells (Jansen *et al*, 2021) and perforin/granzyme B–negative CD56hi NKreg cells (Lauener *et al*, 2023), etc. In the future, by generating “super-tolerant fish” that can simultaneously induce Treg, Breg, and Treg cells, it is expected to achieve efficient xenogeneic organ transplantation. In summary, we present a Treg-induction-based approach in zebrafish that enables efficient immune tolerance and has broad potential applications in transplantation biology, germline engineering, and gene editing.

## Methods

### Ethics statement

All animal experiments were conducted according to the standard animal guidelines approved by the Animal Care Committee of the University of Chinese Academy of Sciences and the Institute of Hydrobiology, Chinese Academy of Sciences.

### Zebrafish maintenance

All the zebrafish lines used in this study were AB and raised in the China Zebrafish Resource Center of the National Aquatic Biological Resource Center (CZRC-NABRC, Wuhan, China, http://zfish.cn). Zebrafish maintained under a 14-h light/10-h dark cycle at 28°C. The embryos were staged according to morphology, as previously described (Kimmel *et al*, 1995). The zebrafish experiments were performed under the approval of the Institutional Animal Care and Use Committee of the Institute of Hydrobiology, Chinese Academy of Sciences under protocol number IHB2014-006. *Tg(CMV:foxp3a)* (ihb946Tg) fish were maintained the same as wild-type fish. Embryos (0-4 dpf) were maintained in 90 mm Petri dishes with 0.3 × Danieau’s buffer. larval stages (5-12 dpf) were transferred to 1 L small round barrel with a filter screen with zebrafish culture water receiving three daily feedings of standardized commercial pellet diet (50 μm). All fish were maintained in recirculating aquaculture systems (RAS) from 12 dpf onward, receiving three daily feedings of either freshly hatched Artemia nauplii (brine shrimp) or standardized commercial pellet diet (100-400 μm).

### Generation and characterization of transgenic fish

The CDSs of the *foxp3a* (NM_001329567.1) was inserted into the vector backbone *Tol2(CMV-sv40; CMV:mCherry)* and purified using DNA extraction reagent (Solarbio). Following microinjection of transgenic plasmid vector (50 ng/µL) and *tol2* mRNA (200 ng/µL) into 1-cell stage *Casper* zebrafish embryos, two sequential rounds of mCherry fluorescence screening were systematically conducted using a Zeiss Axio Imager microscope (AX10) at critical developmental stages: 2 dpf (pre-hatching phase) and 4 dpf (pre-feeding phase). Embryos exhibiting positive fluorescence signals were designated as F0 founders, which were subsequently transferred to standardized zebrafish housing facilities and maintained according to established husbandry protocols.

### Reverse-transcription Quantitative PCR (RT-qPCR)

Total RNA was isolated from the tissues of wildtype and *Tg(CMV:foxp3a)* using TRIzol (Invitrogen). cDNA was synthesized using an oligo-dT primer and RevertAid First Strand cDNA Synthesis Kit (Thermo Fisher Scientific). RT-qPCR was performed using the SYBRGreen Supermix from BioRad (USA) on a BioRad CFX96. The samples were tested in biological triplicates for each gene, and resultant Cq values were averaged. The primers used for RT-qPCR shown in table S2.

### Whole-mount *in situ* hybridization

For whole-mount in situ hybridization (WISH), embryos of 4 dpf were fixed with 4% PFA/PBS overnight at 4°C. WISH was performed with the following probes: *foxp3a*, *CD4-1*, *CD8a*, *ikaros*, *rag1*, *ccr9a*, *cxcr3.2*, the primers used for situ probes vector construction shown in table S1. Generate antisense RNA probes by in vitro transcription using T3/T7/SP6 RNA polymerases with DIG-11-UTP labeling. WISH was performed essentially as described previously: day1-hybridization with probes, day2-incubate with anti-DIG-AP Fab fragments, and day3-develop signal using NBT/BCIP substrate (Roche) (Thisse *et al*, 2004). Embryos hybridized with those probes were mounted in 100% glycerin and acquired of images using a 14× objective (Zeiss, AX10).

### Purification and characterization of male germline stem cells (mGSCs)

Testicular tissues were dissected from six 3-month-old *Tg(ddx4:GFP)* transgenic males and mechanically dissociated using sterile surgical scissors. Spermatogonial purification was performed essentially as described previously (Zhang *et al*., 2022). Purified spermatogonia were resuspended in 600 μL of Leibovitz’ s L-15 medium (Gibco) supplemented with 2% heat-inactivated fetal bovine serum (FBS; Sigma-Aldrich), and loaded into a customized glass capillary tube for intraperitoneal GSC transplantation (IGCT).

### Preparation of sterile host for transplantation

All the WT and *Tg(CMV:foxp3a)* hosts were injected with *dnd1* morpholino (5’-GCTGGGCATCCATGTCTCCGACCAT-3’,50 μM) at 1-cell stage (Weidinger *et al*, 2003), and maintained the hosts to 2 mpf as described (Kimmel *et al*., 1995).

### Intraperitoneal transplantation of purified mGSCs

For the IGCT, the 2 mpf hosts were anaesthetized with 1 × MS-222, and position the hosts in a sponge with the cloacal ventrum oriented dorsally. Intraperitoneal transplantation is administered via the cloacal opening using a glass capillary tube, with 10 μL injected per hosts. After transplantation, all the hosts were examined by a fluorescence microscopy and the potential germline chimeras were identified based on the presence of green fluorescence-positive cells in the abdomen region acquired of images using a 7 × objective (Zeiss, AX10). 30 hosts were anaesthetized after 35 dpt, and the colonization rates of the transplanted spermatogonia as well as its ability to proliferate in host gonads were evaluated.

### Subcutaneous transplantation of trunk tissue of larval fish

The donor of graft was obtained from 4 dpf *Tg(ddx4:GFP)* transgenic founder fish with PGCs GFP fluorescence positive. Gonadal primordia containing primordial germ cells (PGCs) and gonadal somatic cells were microdissected through careful removal of excess tissues by microsurgical knife under stereomicroscopic guidance. The hosts were 2 mpf wildtype and *Tg(CMV:foxp3a)* transgenic fish. Hosts were anesthetized using and a subcutaneous pocket was created in the host’s lateral musculature using a microsurgical knife and fine forceps. Transplantation the graft into the host’s pocket, then transferred the host into 0.3 × Danieau’s buffer supplemented with 1 × antibiotic for recovery. After transplantation, graft colonization status was monitored daily through GFP fluorescence using confocal microscopy. Longitudinal tracking was performed with fluorescence imaging at 7-day intervals until definitive gonadal integration was observed.

### Scanning electron microscope (SEM)

The procedure used to prepare sperm for SEM was described in the previously published article (Zhang *et al*., 2022). Briefly, samples fixed with 2.5% glutaraldehyde overnight at 4°C and dropped onto cell slides and washed with phosphate-buffered saline (PBS, pH7.4) 3 times for 5 min each. After the samples were naturally dried, the samples were dehydrated in a gradient with 30%, 50%, 70%, 100% and 100% ethanol 15 min each. The dried samples were sprayed with gold (Hitachi, E-1010) and used for SEM (Hitachi, S-4800) observation.

### Immunofluorescence on tissue sections and cell suspensions

Immunofluorescence was performed as previously described (Ye *et al*., 2023). Cells and tissue sections of the testis were fixed with 4% paraformaldehyde (PFA) at RT for 4 h for immunofluorescence staining. For the fixed tissue sections were washed with PBST (pH7.4, 0.1% triton-100) 3 times for 5 min each. For the fixed cells, dropped onto adhesive-coated slides and air-dried at room temperature (RT). The cells and tissue sections permeabilized with PBST (pH7.4, 0.5% triton-100) for 30 min, and washed with blocking solution (0.1% triton-100, 2% BSA, and 1% DMSO in PBS) 2 times for 10 min each at RT. The cells and tissue sections were blocked with blocking solution for at least 1 h at RT. The antibodies (1:500 in blocking solution) were used to incubate the samples overnight at 4°C. After washing with blocking solution 3 times for 5 min each, the sections were incubated in secondary antibody (Goat-anti-Rabbit Alexa Fluor 680, 1 mg/mL in stock, 1:1000 in blocking solution) (Thermo Fisher) overnight at 4°C. After washing with PBS 3 times for 5 min each, the sections were stained with DAPI at 1µg/mL at 4°C for 1 h and washing with PBS 3 times for 5 min each, the samples were used for fluorescence microscopy imaging. The following antibodies were used for immunofluorescence staining. Anti-Vasa, anti-Nanos2, anti-Sycp3, and anti-Pcna were against the antigens and purified using antigen-affinity chromatography by our lab. The effectiveness and specificity of antibodies have been described in a previous study (Ye *et al*., 2023).

### RNA sequencing and analysis

For each sample, 200 ng total RNA from the thymus and graft of 10 dpt and 14 dpt of WT or *Tg(CMV:foxp3a)* was used for synthesis and amplification of cDNA, a total of three biological replicates were performed. For RNA-Seq analysis, first, FastqC (v0.12.1) was used to evaluate the raw read quality. Reads were mapped to the corresponding reference genome (GRCz11) using Hisat2 (v2.2.1). The data were converted into BAM format by Hisat2, and BAM files were sorted by Samtools (v1.13). BAM files were counted by featureCounts. Normalization and differential expression analysis were conducted with DESeq2 (v1.48.2) in R, applying a false discovery rate (FDR) cutoff of 0.05. Functional enrichment analysis of differentially expressed genes (DEGs) was performed using clusterProfiler (v3.16.1) against Gene Ontology (GO) and KEGG pathway databases.

### Sperm squeezing and artificial fertilization

Males were anesthetized with 1× MS-222, dried gently, and immobilized ventral-side up on moistened sponge pads. Milt was collected via abdominal pressure application under stereomicroscopic guidance (Leica M205 FA), aspirated from the genital pore using a 10 μL capillary pipette and suspended in 200 μL Hank’s Buffer. The buffer containing milt was wrapped in aluminum foil and maintained on ice until use. Subsequently, 100 μL sperm suspension was added directly on the top of the eggs for in vitro fertilization (IVF), followed by activation with 1 mL system water. Post-fertilization chorion expansion was monitored for 15 min at 28.5°C under controlled incubation conditions (Kimmel *et al*., 1995). For graft-derived sperm collection and artificial fertilization, host fish were anesthetized in 1 × MS-222, and the area surrounding the graft was thoroughly dried. Under a dissecting microscope, 3–5 scales adjacent to the transplantation site were carefully removed with forceps. A superficial incision was then made in the nearby skin using a surgical blade, followed by gentle separation of the epidermis from the graft with forceps. A portion of the whitish testicular tissue was excised, transferred into 200 μL of Hank’s buffer, and homogenized with a microtube pestle to release spermatozoa. Throughout the procedure, the sperm suspension was kept on ice. The resulting sperm solution was immediately mixed with freshly collected eggs for in vitro fertilization (IVF), and fertilization success was evaluated 30 minutes after insemination.

## Data analysis

Early embryos were imaged using a fluorescence stereomicroscope (Axio Zoom.V16, Zeiss), while sections and cells were imaged using a confocal microscope (SP8, Leica). Immunostaining experiments were repeated at least three times and representative example are shown. The unpaired two-tailed Stugent’s t-test was used to calculate the P values. All data are presented as mean ± standard deviation (mean ± SD). GraphPad Prism 8.3.0 was used for graph plotting and statistical analysis.

## Data availability

The data that support the findings of this study are available in the article and supplementary material of this article.

## Acknowledgments

We sincerely thank Luyuan Pan and Linglu Li from the China Zebrafish Resource Center (CZRC) for their technical assistance *in vitro* fertilization, Fang Zhou and Guangxin Wang from the analytical and testing center of Institute of Hydrobiology, CAS for their support with confocal imaging, and Yuan Xiao and Zhengfei Xing from the analytical and testing center of Institute of Hydrobiology, CAS for their assistance with sperm imaging. This work was supported by the National Natural Science Foundation of China (32025037 and 32403020), the Strategic Priority Research Program of Chinese Academy of Sciences (CAS) (XDB0730300), the STI2030—Major Projects (2023ZD04055), the National Key R&D Program of China (2023YFD2400200), Natural Science Foundation of Hubei Province (2025AFA053), Natural Science Foundation of Wuhan (2024040701010069), Science and Technology Special Fund of Hainan Province (ZDYF2024XDNY256), and State Key Laboratory of Breeding Biotechnology and Sustainable Aquaculture (2024BBSA01).

